# Measuring Protein Binding to Lipid Vesicles by Fluorescence Cross-Correlation Spectroscopy

**DOI:** 10.1101/146464

**Authors:** Daniela Krüger, Jan Ebenhan, Kirsten Bacia

## Abstract

Fluorescence Correlation Spectroscopy (FCS) has been previously used to investigate peptide and protein binding to lipid membranes, as it allows for very low amounts of sample, short measurement times and equilibrium binding conditions. Labeling only one of the binding partners however comes with certain drawbacks, as it relies on identifying binding events by a change in diffusion coefficient. Since peptide and protein aggregation can obscure specific binding and since non-stoichiometric binding necessitates the explicit choice of a statistical distribution for the number of bound ligands, we additionally label the liposomes and perform dual-color Fluorescence Cross-Correlation Spectroscopy (dcFCCS). We develop a theoretical framework showing that dcFCCS amplitudes allow calculation of the degree of ligand binding and the concentration of unbound ligand, leading to a binding model-independent binding curve. As the degree of labeling of the ligands does not factor into the measured quantities, it is permissible to mix labeled and unlabeled ligand, thereby extending the range of usable protein concentrations and accessible dissociation constants *K_D_*. The total protein concentration, but not the fraction of labeled protein needs to be known. In this work, we apply our dcFCCS analysis scheme to Sar1p, a protein of the COPII complex, which binds ‘major-minor-mix’ liposomes. A Langmuir isotherm model yields *K_D_* = (2.1±1.1)*μM* as the single site dissociation constant. The dual-color FCCS framework presented here is highly versatile for biophysical analysis of binding interactions. It may be applied to many types of fluorescently labeled ligands and small diffusing particles, including nanodiscs and liposomes containing membrane protein receptors.

## Introduction

Quantitative analysis of protein binding to lipid membranes is an essential technique both for understanding native biological processes and for pharmaceutical drug development. A typical model system for investigating interactions between proteins and lipids as well as interactions between any type of ligands and membrane proteins is provided by membrane vesicles with diameters on the order of 100 nm. Membrane extrusion and detergent removal are commonly used techniques for producing mostly unilamellar vesicles with relatively low size dispersion.

Although a number of techniques is available for quantifying binding of proteins or other ligands to membrane vesicles, each has its limitations, calling for an analytical method that allows quantitative analysis of binding with the following features: (I) measurement under equilibrium conditions, (II) freely diffusing particles, no interference from a support, (III) output of titration curves with no requirement for a pre-defined binding model, (IV) low sample consumption.

Fluorescence Correlation Spectroscopy (FCS) is a low-invasive, single-molecule sensitive fluctuation technique that was first introduced in the 1970s (1). Benefitting from technological advances, it became more widely applicable in the 1990s (2) and has by now matured into a versatile and robust technique that is widely applicable to biological macromolecules and assemblies, including proteins, nucleic acids and lipid assemblies (3, 4). FCS and its related fluorescence fluctuation techniques are highly specific and sensitive, permitting analysis of interactions even in complex systems including membrane preparations and living cells. From a practical point of view, it is advantageous that only microliters of material and acquisition times on the order of seconds to minutes are needed. Moreover, commercial setups are available for confocal microscope-based FCS.

Protein-membrane interactions have been investigated in two basic geometries: the planar membrane configuration (Fig. 1 A), and the small diffusing particle configuration (Fig. 1 C). In the first geometry, a planar membrane intersects the confocal detection volume. Giant unilamellar vesicle bilayers, supported bilayers and lipid monolayers can be used as effectively planar membranes. The lateral diffusion of fluorescently labeled, membrane-bound protein in the plane of the membrane is detected, along with the 3-dimensional diffusion of free protein in one half of the detection volume. The diffusion of a membrane-bound protein is significantly slower than diffusion of the free protein in aqueous solution, allowing for two-component fitting (see e.g. (5, 6)). However, the detected fraction of free protein is sensitive to membrane positioning. Analogous binding experiments have been performed with small fluorescent ligands binding to membrane proteins (7).

**Figure 1.**
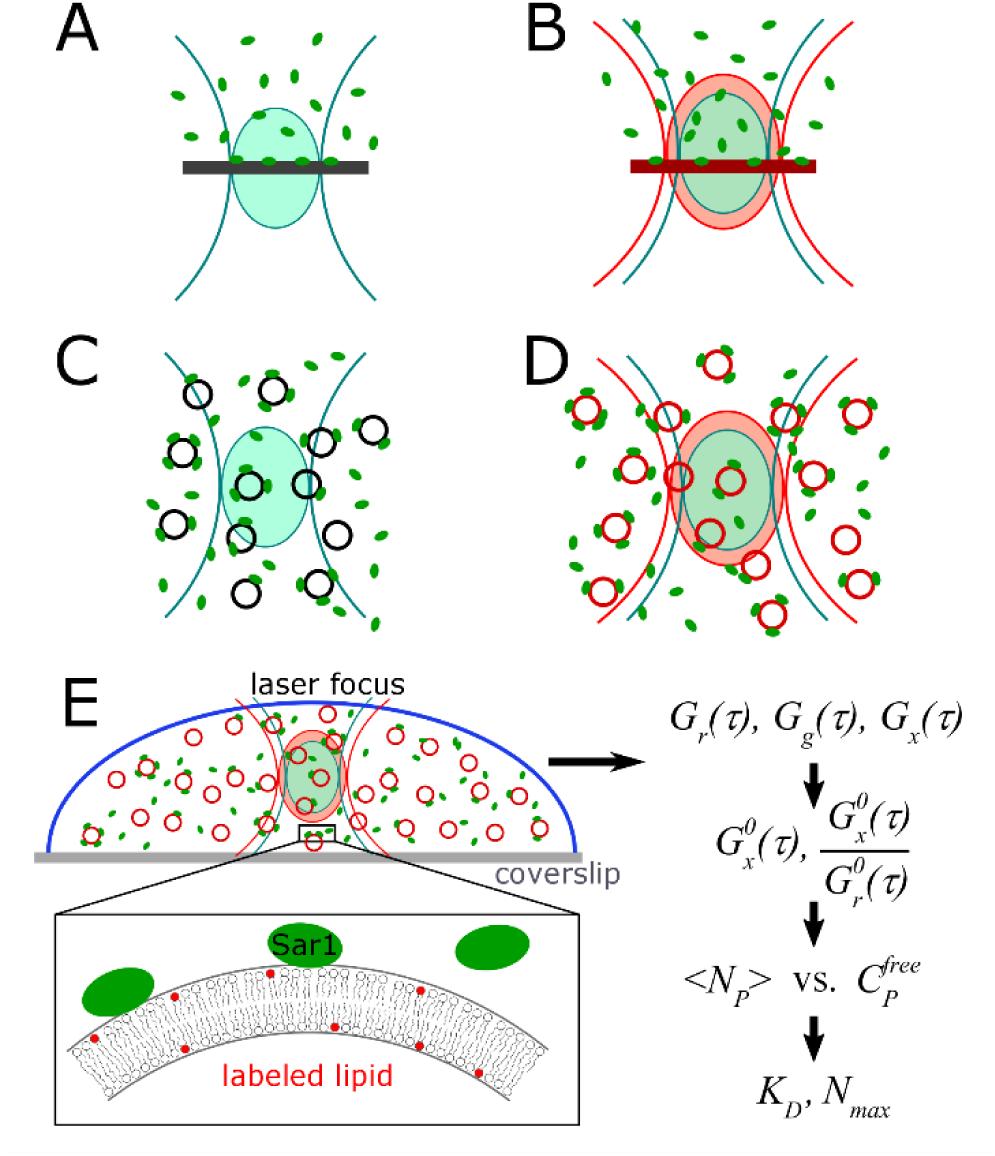
FCS/dcFCCS configurations for analyzing ligand-membrane binding. (A) Autocorrelation analysis of a labeled ligand binding to an unlabeled planar membrane. Fractions of bound and free ligand are quantitated by a two-component diffusion model. Measured fractions are sensitive to focus position. (B) Same geometry as in (A), with additional exploitation of a distinctly labeled binding partner in the membrane and dual-color cross-correlation. (C) Analysis of freely diffusing ligand-liposome particles. The ligand is labeled; free and liposome-bound fractions are quantitated by a two-component diffusion model. Influence of binding stoichiometry needs to be considered or liposomes used in large excess (8-10). (D) Same geometry as in (C), with additional exploitation of distinctly labeled liposomes. (E) Implementation of configuration (D) in this work. The dcFCCS foci are placed on the binding reaction, which occurs in an aqueous buffer. The liposome bilayer is labeled with a red-fluorescent lipid analog. Green-fluorescently labeled Sar1p protein constitutes the ligand that binds to the liposomes. Using the cross-correlation and the red autocorrelation amplitude, the degree of ligand binding vs. free ligand is obtained. A Langmuir isotherm model is applied, yielding the number of total binding sites and the dissociation constant.

The second configuration uses small protein-lipid particles that diffuse through the confocal detection volume as whole entities, namely small liposomes, proteoliposomes, micelles or lipid nanodiscs (Fig. 1C). Again, the different diffusion coefficients of free and bound protein allow to monitor binding. In this way, binding of fluorescently-labeled protein to unlabeled liposomes (8-10) and nanodiscs has been analyzed (11). All these applications relied on single color detection of the protein and extracted the unbound and bound fractions from two-component diffusion models.

In complex systems that contain a variety of potential interaction partners for the ligand of interest, diffusion analysis alone is insufficient for pinpointing molecular interactions, as any of the potential interactions may lead to slower diffusion. Dual-color Fluorescence Cross-Correlation (dcFCCS) (12) circumvents this specificity problem by labeling the potentially interacting binding partners with spectrally distinguishable dyes and monitoring their dynamic co-localization. If the binding stoichiometry is not 1:1, brightness distributions need to be accounted for (13). General advantages of dcFCCS analysis for binding analysis are provided by the increased specificity and the robustness in obtaining amplitudes as opposed to fractions in two-component fits.

In the planar membrane configuration (Fig. 1B), dcFCCS has been applied to monitor the binding of a ligand to a specific membrane constituent as well as to the in-plane association of membrane constituents (6, 14, 15).

In the small particle configuration (Fig. 1D), dcFCCS has been used to monitor co-localization of protein cargo in endocytic vesicles (5), vesicle docking (16) and reconstitution of a membrane protein into liposomes (17). However, to our knowledge, protein binding to liposomes has so far not been analyzed truly quantitatively by using dcFCCS titration curves. In this paper, we derive the framework for obtaining titration curves (degree of binding vs. free ligand) from dcFCCS data and quantitate the binding of the small GTPase protein Sar1p to liposomes. We show that, in addition to the general dcFCCS advantages, a number of assumptions made in single color FCS analysis (9) are no longer required when using dcFCCS. Furthermore we demonstrate that typical challenges in dcFCCS analysis, namely a need for stoichiometric protein labeling and a limited range of usable concentrations do not apply here.

The Sar1 protein is a small GTPase of the Sar1/Arf1 family, which belongs to the superfamily of Ras GTPases (18, 19). Upon activation by GTP, Sar1 binds to the Endoplasmic Reticulum membrane in eukaryotic cells by embedding an N-terminal amphipathic helix into the proximal (i.e., cytosolic) leaflet of the bilayer (28). Activated Sar1p is implicated in membrane curvature generation. Furthermore, it initiates the recruitment of further COPII coat components, leading to the formation of a COPII-coated bud and, ultimately, intracellular transport vesicles. The COPII transport machinery is ubiquitous among eukaryotes; here we study the Sar1 protein (Sar1p) from baker’s yeast (*Saccharomyces cerevisiae*), which is permanently activated by the non-hydrolyzable GTP-analog GMP-PNP.

## Theory

A comprehensive review of FCS including dcFCCS can be found in (4). Here we demonstrate a theoretical framework for analyzing dcFCCS data of protein-liposome binding titrations, where the protein (or other ligand) and liposome are labeled with spectrally distinct fluorophores.

For simplicity only, the protein-label will be referred to as “green” and the liposome-label as “red” (see Table 1 for a list of variables). To begin with, it is assumed that each protein molecule carries exactly one green fluorophore, but it will be shown further below (Eq. 12) that this is not a requirement. Liposomes are labeled with a small fraction of red lipid analog, resulting in a Poisson distribution of the dye molecules among the lipids.

**Table 1.** List of variables

| Variable | Meaning |
| --- | --- |
| index $g$ | pertaining to the green detection channel |
| index $r$ | pertaining to the red detection channel |
| index $x$ | pertaining to the cross-correlation channel |
| $F_a(t)$, where $a = g, r$ | fluorescence intensity in channel $a$ |
| $B_a$, where $a = g, r$ | mean background count-rate in channel $a$ |
| $G_a(\tau)$, where $a = g, r, x$ | correlation functions |
| $\tau$ | correlation time |
| $G_{a,m}^0$, where $a = g, r, x$ | measured correlation amplitudes |
| $G_a^0$, where $a = g, r, x$ | corrected correlation amplitudes |
| $\tau_D$ | diffusion time |
| $D$ | diffusion coefficient |
| $\tau_T$ | blinking relaxation time |
| $T$ | average fraction of dye in the dark state |
| $\omega_{xy}$ | lateral focus radius |
| $\omega_z$ | axial focus radius |
| $S$ | structure parameter |
| $V_{eff,a}$, $a = g, r, x$ | effective focal volume of channel $a$ |
| $f_f$ | fractional amplitudes of the free diffusing protein |
| $f_v$ | fractional amplitudes of the liposome-bound protein |
| $N_g$ | number of green fluorophores on a vesicle |
| $C_g^0$ | total concentration of labeled protein |
| $C_g^{free}$ | concentration of free labeled protein |
| $\chi_g$ | labeling fraction |
| $N_p$ | number of proteins on a vesicle |
| $C_p^0$ | total concentration of protein |
| $C_p^{free}$ | concentration of free protein |
| $C_{sites}^{free}$ | concentration of free binding sites |
| $C_{sites}^{bound}$ | concentration of occupied binding sites |
| $K_D$ | dissociation constant |
| $N_{max}$ | number of binding sites per vesicle |
| $N_A$ | Avogadro constant |

The measured cross-correlation between red and green channel *G_x_*(*τ*) and the measured autocorrelation in the green channel are given by

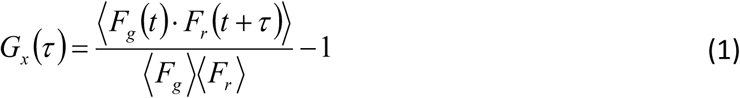

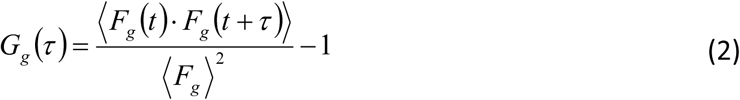

Here, 〈…〉 denotes a time average over the acquisition time. The red channel autocorrelation, *G_r_*(*τ*), is treated in analogy to the green.

The autocorrelation function obtained in the red channel stems from the three-dimensional diffusion of the red-labeled liposomes. It is fitted with a model equation that contains one blinking term (*τ_T_*: blinking relaxation time, *T*: average fraction of dye in the dark state) and a diffusion term (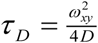: diffusion time, 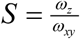: structure parameter) (4):

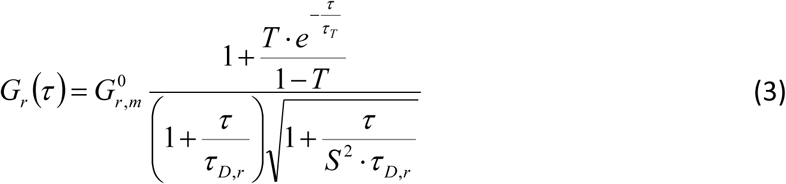

The green channel autocorrelation (corresponding to the labeled protein) has to be fitted with two components, where *f_v_* and *f_f_* denote the fractional amplitudes of the liposome-bound and the free diffusing protein, respectively (4):

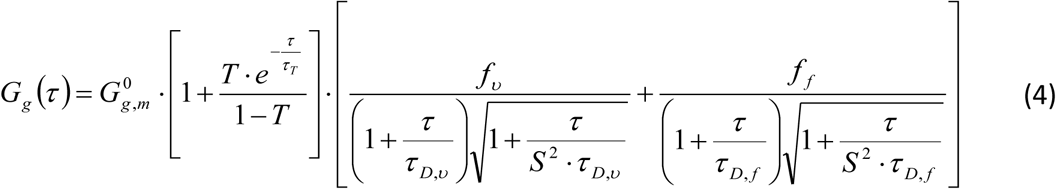

The cross-correlation includes only the vesicle component and no blinking:

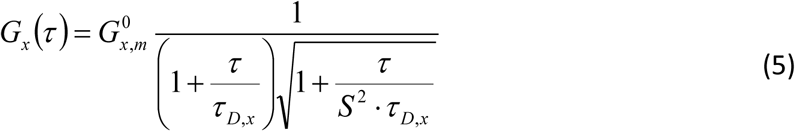

Measured autocorrelation amplitudes (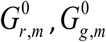) are corrected for uncorrelated background (20) by multiplying them by 〈*F_a_*〉^2^ /(〈*F_a_*〉−〈*B_a_*〉^2^), where 〈*F_a_*〉 is the mean total count-rate and 〈*B_a_*〉 is the mean background count-rate in the respective channel *a*. The measured cross-correlation amplitude 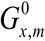 is background-corrected by multiplying by 〈*F_g_*〉〈*F_r_*〉 /[(〈*F_g_*〉 −〈*B_g_*〉)(〈*F_r_*〉−〈*B_r_*〉)]. The red autocorrelation amplitude and the cross-correlation amplitude are then cross-talk corrected according to (21).

To analyze binding of proteins to liposomes by dcFCCS, only the corrected autocorrelation amplitude of the red channel 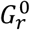 as well as the corrected cross-correlation amplitude 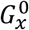 are needed. (In addition, *τ_D_* from the red autocorrelation curve is used to check vesicle size.)

Taking into account the various particle brightnesses (4, 13) and disregarding potential quenching of the green-labeled protein upon binding to the liposomes, the amplitudes are given by the following equations:

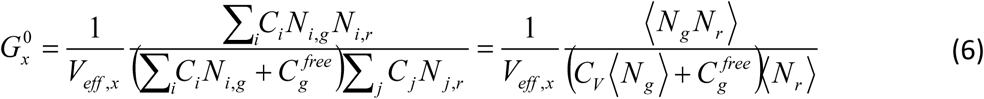

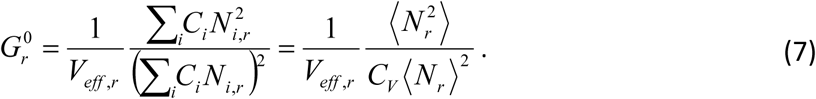

Here, the angle brackets 〈*…*〉 are to be understood as an ensemble average. *N_i,g_* denotes the number of green fluorophores (labeled proteins) on a vesicle of species *í* with concentration *C_i_*. Likewise *N_i,r_* stands for the number of red fluorophores (labeled lipids). The concentration *C_v_* is the total concentration of vesicles, regardless of the number of fluorophores attached to them. All concentrations are to be understood as “particles per volume” and can be converted into molar quantities by division by Avogadro’s constant: *c* = *C / N_A_*.

The effective focal volumes *V_eff,x_* and *V_eff,r_* are determined by calibration measurements (see Materials and Methods section).

The total protein concentration is the sum of the free protein concentration and the sum over all protein bound to vesicles:

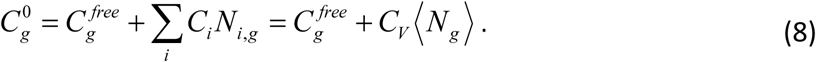

We assume that the binding of proteins is not influenced by the number of lipid dye molecules per vesicle. In that case, *N_g_* and *N_r_* are stochastically independent variables and〈*N_g_N_r_*〉=〈*N_g_*〉〈*N_r_*〉.

We further assume a Poisson distribution for the number of lipid dye molecules *N_r_*. The variance of a Poisson distribution equals its mean: 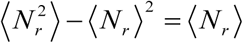.

Hence, the correlation amplitudes simplify considerably:

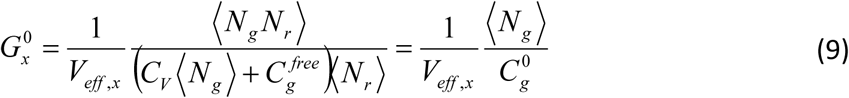

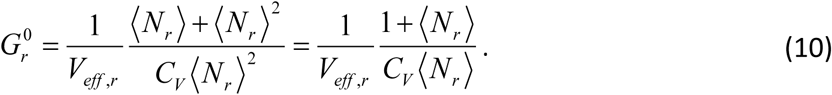

From the relative cross-correlation,

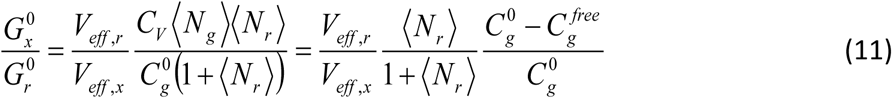

the concentration of free protein 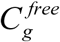 can be calculated, provided that the total protein concentration 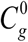 is known. The term 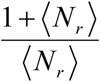 becomes negligible at sufficient lipid dye concentrations, as shown in the Results section.

Since the cross-correlation amplitudes become small at high protein concentrations, a mixture of labeled and unlabeled protein is used for these samples with a total concentration of 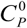 and a labeling fraction *χ_g_*. This only changes 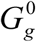, which is not used in this work, whereas 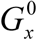 is unaffected. This can be seen as follows:

The total concentration of labeled protein is given by 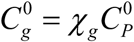. We assume that the labeling does not alter the binding properties of the protein, in which case the concentration of unbound labeled proteins is given by 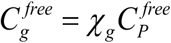. The number of labeled protein molecules *N_g_* among all protein molecules bound to a liposome (regardless of a label) *N_p_* follows a binomial distribution *N_g_* ~ *B*(*N_p_*, *χ_g_*) yielding:

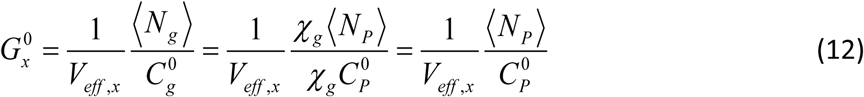

Therefore, it is permissible to concomitantly replace 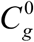 by 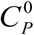, 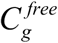 by 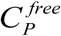 and 〈*N_g_*〉.

As a side note, because the equations for 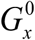 and 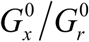 contain only mean numbers of green fluorophores, even though this work uses protein molecules that have been labeled at a single engineered cysteine residue, protein preparations that are stochastically labeled at multiple sites per protein can also be used.

We are now able to determine the degree of protein binding from the absolute cross-correlation amplitude as

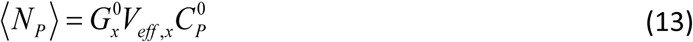

and the free protein concentration 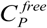 from the relative cross-correlation as

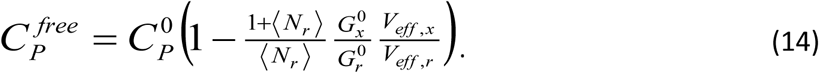

Plotting 〈*N_p_*〉 versus 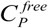 yields a standard, binding model-independent ligand binding curve (22).

The half-saturation point of this curve allows for a first estimation of the affinity of the protein to liposomes.

The ligand binding curve ( 〈*N_p_*〉 vs. 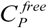) is the basis for testing any specific binding model (22).

Here we use the model of *N_max_* identical and independent binding sites (non-cooperative binding, (22)), which is also referred to as the Langmuir isotherm:

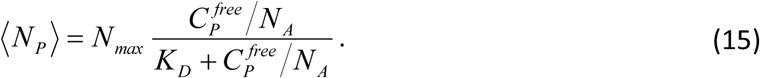

The dissociation constant *K_D_* indicates the molar concentration of free protein, at which half of the binding sites are occupied.

## Materials and Methods

### Liposomes

Liposomes were prepared from a complex mixture of lipids (termed the ‘major-minor-mix’) that was previously established in COPII *in vitro* reconstitution experiments (23-25). It consists of 34.4 mol% of 1,2-dioleoyl-*sn*-glycero-3-phosphocholine (DOPC), 14.8 mol% of 1,2-dioleoyl-*sn*-glycero-3- phosphoethanolamine (DOPE), 3.4 mol% of 1,2-dioleoyl-*sn*-glycero-3-phosphate (DOPA), 5.4 mol% of L-α-phosphatidylinositol from soy (soy-PI), 1.5 mol% of L-α-phosphatidylinositol-4-phosphate from porcine brain (PI(4)P), 0.5 mol% of L-α-phosphatidylinositol-4,5-bisphosphate from porcine brain (PI(4,5)P2), 1.3 mol% of 1,2-dioleoyl-*sn*-glycero-3-(cytidine diphosphate) (CDP-DAG) and 33.3 mol% of ergosterol. All phospholipids were purchased from Avanti Polar Lipids, Alabaster, AL), ergosterol was from (Sigma-Aldrich, Germany). Lipids were dissolved in organic solvent (chloroform:methanol 2:1 (vol/vol)) and doped with 0.004 mol% of the red fluorescent lipid analog DiIC18(5) (DiD) (1,1’- dioctadecyl-3,3,3’,3’-tetramethylindodicarbocyanine, 4-chlorobenzenesulfonate, from Invitrogen/Thermo-Fisher, Carlsbad, CA). Solvents were removed under vacuum in a rotary evaporator to obtain a thin lipid film. The lipid film was subsequently hydrated with aqueous HKM buffer (20 mM HEPES/potassium hydroxide, pH 6.8, 50 mM potassium acetate and 1.2 mM magnesium chloride) for 1 hour at a total lipid concentration of 4 mM. Subsequently, the lipid suspension was extruded through a 50 nm polycarbonate membrane using an Avanti Mini Extruder (both obtained from Avanti Polar Lipids, Alabaster, AL). The size of the liposomes was measured by dynamic light scattering (DLS) (Zetasizer Nano S, Malvern Instruments, UK). Lipid concentration was determined using a malachite green phosphate assay. Liposomes were stored at 4°C for a maximum of one week prior to dcFCCS measurements.

### Protein binding

The protein Sar1p from *Saccharomyces cerevisiae* was expressed and purified as previously described (24). The protein variant Sar1pS147C/C171S was prepared in the same way, but additionally labeled with Alexa Fluor 488 maleimide (Invitrogen/Thermo-Fisher, Calsbad, CA). Both labeled and unlabeled Sar1p proteins were functional in a COPII recruitment and membrane deformation assay (24) and with respect to GTPase enzyme activity. Protein concentrations were determined by UV absorption, using linear unmixing of the protein spectrum and the nucleotide spectrum. Protein concentrations were additionally checked by a Bradford assay. For each series of dcFCCS measurements, a set of eleven to sixteen binding reactions was incubated in parallel. Each sample contained 4 μl HKM buffer, 1 mM GMP-PNP (guanosine 5′-[β,γ-imido]triphosphate, from Sigma-Aldrich, Germany), extruded ‘major-minor-mix’ liposomes at 0.09 mM total lipid concentration and variable total concentrations of Sar1p protein 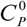, ranging from around 20 nM to 200 μM, with labeling fractions ranging between 100% to 0.08 mol%. Various labeling fractions were used within overlapping concentration regimes.

### Dual-color FCCS and data analysis

DcFCCS measurements were carried out on a commercial inverted confocal fluorescence microscope with a fluorescence correlation spectroscopy unit (LSM710/ConfoCor3 from Zeiss, Jena, Germany). The laser powers were attenuated using an acousto-optical tunable filter to 1.1 to 5.7 μW (488 nm) and 0.8 to 5.5 μW (633 nm, all values behind the objective) for optimal count-rates while avoiding photobleaching. The coverslip was placed directly on the immersion water on top of a C-Apochromat 40x, NA 1.2 objective with correction collar. The dichroic 488/561/633 nm was chosen as the main beam splitter, a longpass 635 nm as the secondary beam splitter. A bandpass emission filter (505 to 610 nm) collected the signal from the green dye and a longpass filter (> 650 nm) the red signal. Avalanche photodiodes served as detectors. Acquisition time was 5 times 1 minute. Auto- and cross-correlation were performed using the ZEN 2009 software (Zeiss, Jena, Germany), which was also used for fitting the FCCS data. Correlation amplitudes from the different samples were used to calculate binding curves, which were fitted with the Langmuir isotherm model in MATLAB (MathWorks, Natick, MA, USA).

### Calibration measurements

The red autocorrelation and the cross-correlation amplitude were corrected for background (20) and for the small cross-talk of the green dye to the red channel (21). The bleed-through ratio of the green-label Sar1p to the red channel was determined in a calibration measurement to be *κ_Gr_* = 0.0087. For the calibration of the detection volumes, Alexa Fluor 488 hydrazide and Alexa Fluor 633 hydrazide (Life technologies, Carlsbad, CA, USA) were used. The structure parameter *S* was determined to be 6.7 ± 0.8 in the green channel and 6.3 ± 0.7 in the red channel. For simplicity, an average structure parameters of 6.5 was used for both channels. The effective detection volume size *V_eff,g_* was calculated from the diffusion time of Alexa Fluor 488 and its known diffusion coefficient (*D_A488_* = 435 *μm*^2^ /*s* (26)), yielding 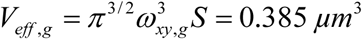. The ratio of the focus radii in the red and the green channel *ω_xy,r_*/*ω_xy,g_* was determined using double labeled vesicles. The calibrations yielded 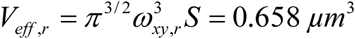 and 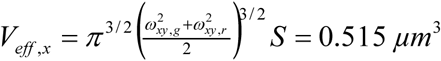.

## Results

We wanted to explore the capabilities of the dcFCCS for quantitating protein-lipid binding. To do so, we chose the yeast protein Sar1p in its GMP-PNP activated, binding-competent state and extruded liposomes. Sar1p was fluorescently labeled in green with Alexa Fluor 488. Extruded liposomes consisted of a defined lipid composition that had previously been established in biochemical experiments (‘major-minor mix’ (24, 25), see Materials and Methods). Liposomes were fluorescently labeled in the far red by 0.004 mol% DiD.

### Test for equilibrium

We first explored the temporal evolution of the binding reaction on the time scale of hours by performing dcFCCS measurements every 15 minutes and monitoring the correlation amplitudes. This pilot experiment was performed on five samples with different protein and lipid concentrations. In all cases, the relative cross-correlation amplitude 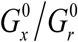 showed no systematic changes after incubation times between 1 and 4 hours (see Fig. 2A for an example). The two-hour timepoint, at which the system appears to be safely at equilibrium, was chosen for all further experiments.

**Figure 2.**
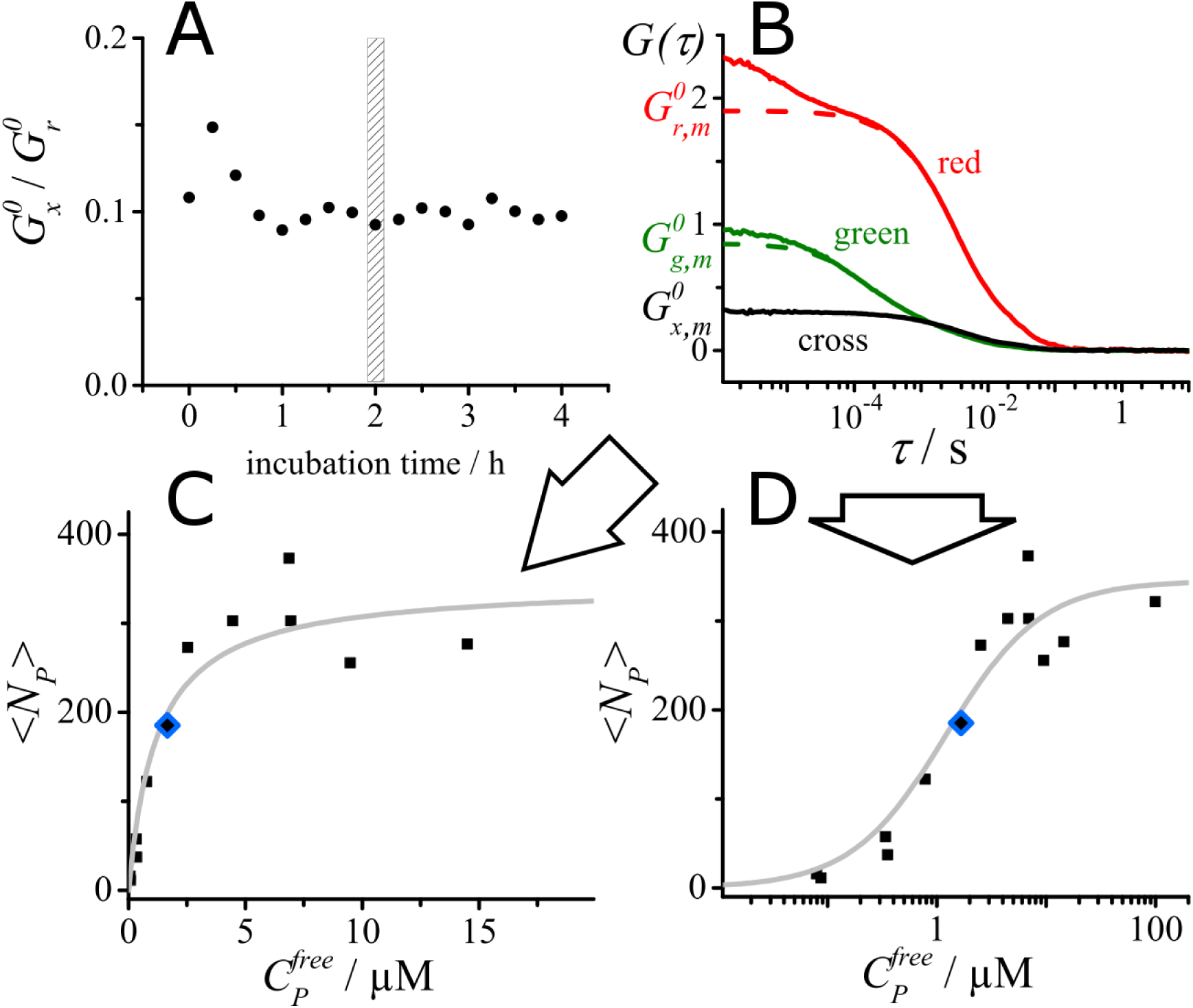
Principle of dual-color FCCS analysis of binding reactions. (A) Relative cross-correlation amplitude from dcFCCS measurements at different time points for a binding reaction consisting of 1 μM Sar1p, 0.2 mM total lipid (‘major-minor mix’ liposomes) and an excess of GMP-PNP (1 mM). The system appears to be equilibrated by the two-hour time point. (B) Dual-color FCCS data analysis of a single protein-lipid binding reaction at the two-hour incubation point (2 μM Sar1p, 0.2 mM total lipid, 1 mM GMP-PNP). The red autocorrelation curve was fit with the diffusion-reaction model (Eq. 3). The dashed curve represents a simulated FCS curve based on the fit parameters, simulated without the blinking term. It indicates the value of the red autocorrelation amplitude, 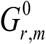. The green autocorrelation curve was fit according to Eq. 4. The dashed curve was again simulated based on the fit parameters but without blinking. It indicates the green autocorrelation amplitude 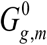. The cross-correlation curve was fit according to Eq. 5. to obtain the cross-correlation amplitude 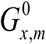. Correlation amplitudes were corrected as described in the Theory and Materials and Methods sections and used together with the total protein concentration to calculate the degree of binding 〈*N_P_*〉 and the concentration of free protein 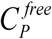 in this sample. (C) Eleven binding reactions were incubated separately, analyzed by dcFCCS and the degree of binding 〈*N_P_*〉 was plotted against the concentration of free protein 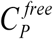. The solid gray line indicates the best fit of the Langmuir isotherm model (independent sites, non-cooperative binding) (Eq. 13). (D) Same plot as in (C) but with 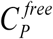 on a logarithmic scale. In this representation, the Langmuir model yields a sigmoidal curve. The blue diamonds in (C) and (D) mark the data point derived from the single binding reaction in panel (B).

### Ligand binding curves from dcFCCS

Figure 2B shows an example of a dcFCCS measurement obtained on a single binding reaction at the two-hour timepoint. Correlation curves were fit according to Eqs. 3 to 5. All amplitudes 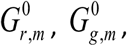, 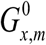 were corrected for minor background count-rates and the red autocorrelation and the cross-correlation amplitude were additionally cross-talk corrected (21), yielding 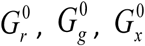. The total protein concentration 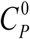 in the reaction was independently determined by a UV absorption assay prior to mixing the components of the binding reaction. Using this total protein concentration as additional information, the degree of protein binding 〈*N_P_*〉 (i.e., the average number of protein molecules per liposome) and the concentration of free (i.e., unbound) protein 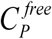 were calculated according to Eq. 13 and Eq. 14, respectively. Detection volumes *V_eff,g_*, *V_eff,r_*, *V_eff,x_* were determined from calibration measurements (see Materials and Methods). The term 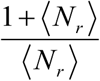 will be discussed further below. This procedure was performed for each of the eleven parallel binding reactions, which contained different concentrations of total protein. The degree of binding 〈*N_P_*〉 was plotted against the concentration of free protein 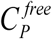, yielding a ligand binding curve (see Fig. 2C for linear axes and Fig. 2D for a semi-logarithmic representation). Here, we chose the simple model of non-cooperative binding of the protein to a number (*N_max_*) of equal, independent binding sites (Langmuir isotherm model).

### Mixing labeled and unlabeled protein

Equation 12 shows that the degree of green labeling of the protein preparation has no influence on the cross-correlation amplitude. Therefore, labeled and unlabeled protein can be mixed at any ratio, which allowed us to span several orders of magnitude of total protein concentration (from 20 nM to 200 μM), without compromising on suitable numbers of fluorescent particles in the focal volume nor on suitable count-rates.

The titration experiment from Figure 2 C/D was repeated using the same liposome preparation (Fig. 3B). In addition, four more ligand binding curves were acquired using independent liposome preparations and independent determinations of protein concentration (Fig. 3 C-F).

**Figure 3.**
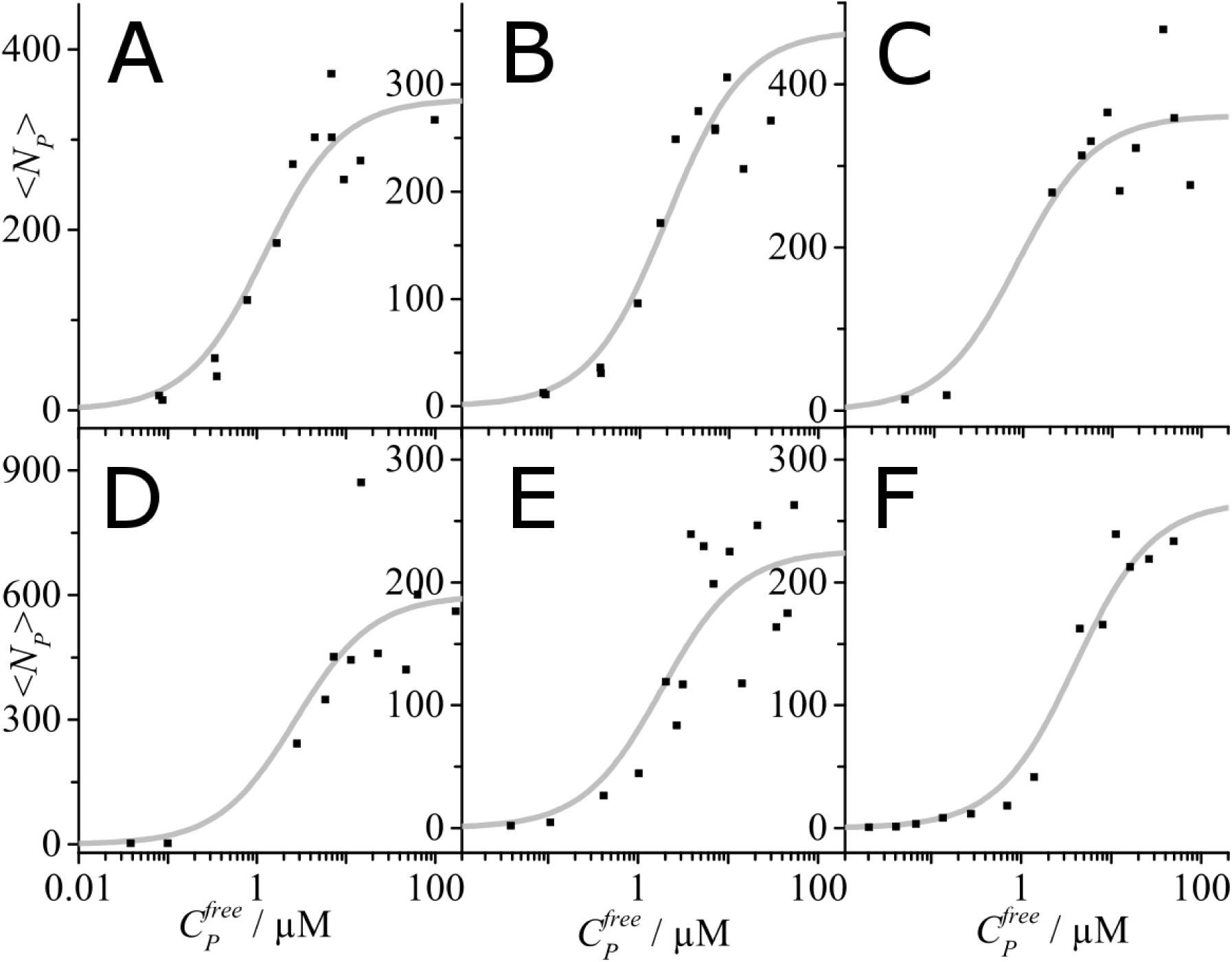
Protein titration curves derived from dcFCCS measurements. In addition to the ligand binding curve already shown in Fig. 2D, five more binding curves were obtained. Each binding curve was modeled with a Langmuir isotherm (gray line).

Taking into account all measurements, various labeling fractions were chosen in each concentration regime as an additional control for any label-induced binding artifacts. For a given protein concentration, the largest and the lowest labeling fractions differed at least by a factor 10, with good overlaps, except for the very low concentration conditions (below 100 nM), where purely labeled protein was required to ensure sufficient signal. No influence of the fluorescent label was apparent.

### Ligand-binding curves

Considering the six titration experiments, average liposome size after extrusion differed slightly between the five liposome preparations, as determined from the FCS diffusion time of the liposomes prior to protein addition (between 79 and 88 nm) and from dynamic light scattering (z-average between 82 and 93 nm; intensity-weighted average between 92 and 100 nm). All six protein titration curves were fit with the Langmuir model. Each resulted in a single binding site *K_D_* in the low μM range, with an average over all binding curves (panels 3A – F) of 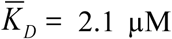 = 2.1 NM and a standard deviation of 1.1 μM. The total number of binding sites was more variable between the different liposome preparations with values of *N_max_* ranging between approximately 200 and 600. The variations in the number of total binding sites is only partially accounted for by different liposome sizes and might actually reflect morphological variations of the liposomes upon binding of the amphipathic helix protein Sar1p, whereby tubular membrane sections are expected to accommodate particularly dense, helical packing of the Sar1 protein (27, 28). Nonetheless, a more complex binding model that involves binding sites of different affinities or cooperativity does not seem justified in this case (Fig. 3 A-F).

### Liposome brightness considerations

When FCCS binding analysis is carried out using the cross-correlation and the red autocorrelation amplitude, the degree of labeling of the green binding partner does not influence the results, but the degree of labeling of the red binding partner does. Nonetheless, uniform brightness of the red-labeled liposomes is not a necessity. This is because stochastic liposome labeling with a small fraction of lipidic dye follows a Poisson distribution, which is easily taken into account and simply yields the pre-factor 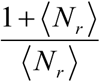 in Eqs. 10 and 11. 〈*N_p_*〉 is the mean number of red fluorescent lipids per liposome. If the liposomes are prepared from lipids with a known area per lipid (determined by x-ray diffraction, (29, 30)) and the diameter is known, it is straight-forward to calculate the average number of fluorescent lipids per liposome from the molar fraction of labeled lipid among all lipids. In the present case however, since the canonical ‘major-minor-mix’ used in COPII studies includes the sterol ergosterol in a complex mixture of lipids (31), the mean area per lipid can only be roughly estimated and there is some uncertainty as to the size distribution and shape of the liposomes.

Nonetheless, the pre-factor 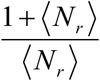 does not constitute a major source of error in the binding analysis, as can be seen from Fig. 4, which shows the best fit {*K_D_*; *N_max_*}-pairs for a range of 〈*N_r_*〉 - values. Assuming a typical mean area per lipid around *A_L_* ≈ 0.6 *nm*^2^ (29) and the diameter of the liposomes to be between 80 nm (from the FCS diffusion time) and 100 nm (from intensity-weighted dynamic light scattering), the *K_D_* varies only by 1.5% and *N_max_* by 0.1%. Therefore, a precise knowledge of 〈*N_r_*〉 is not necessary. *A_L_* ≈ 0.6 *nm*^2^ is an upper bound, because the presence of the small sterol molecule reduces the mean area per lipid, pushing 〈*N_r_*〉 to greater values and thereby reducing its influence on *K_D_* and *N_max_* even further. Generally, choosing a sufficiently high content of labeled lipid renders the influence of the stochastic labeling negligible.

**Figure 4.**
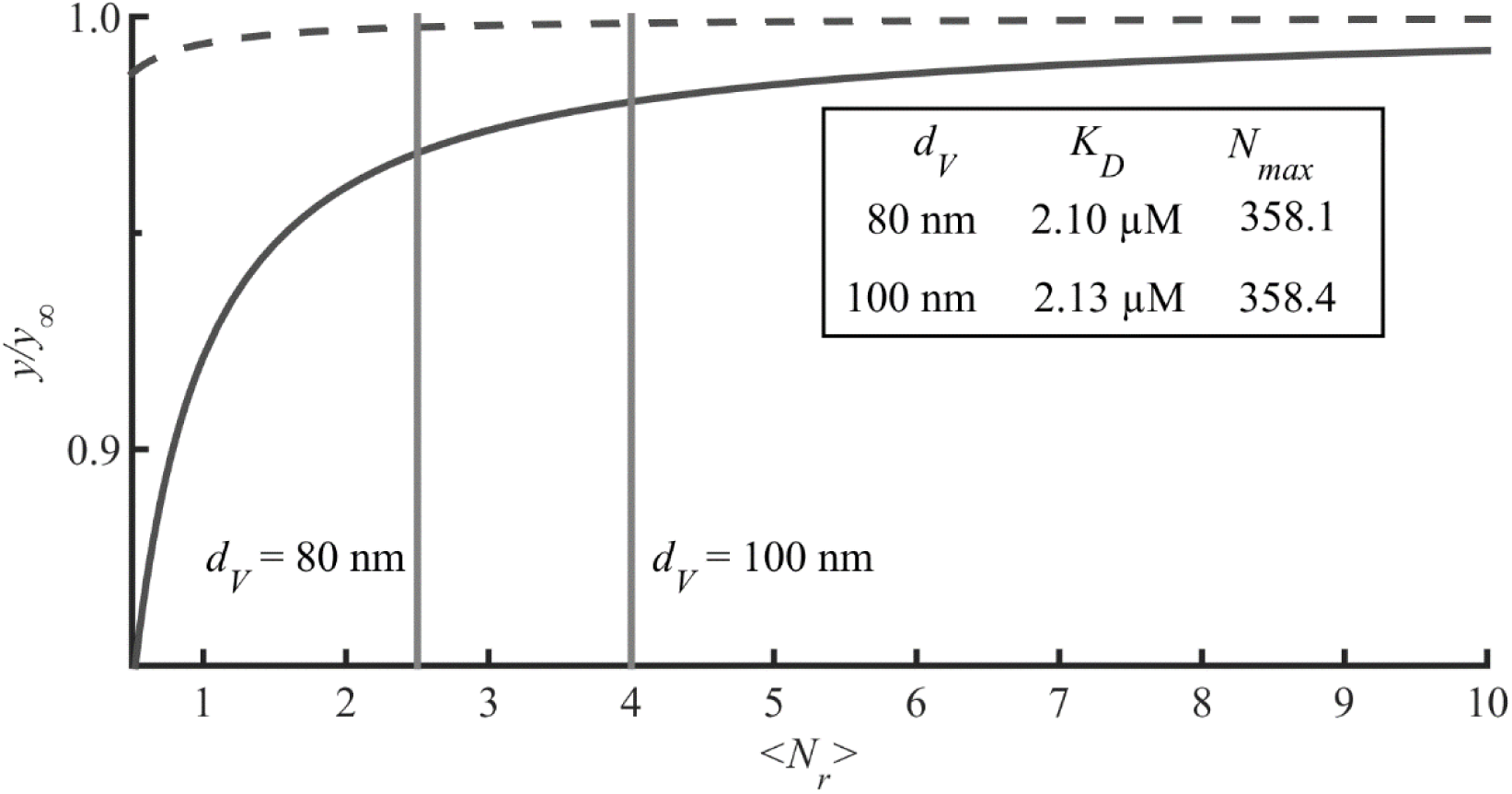
Mean fitted values for *K_D_* and *N_max_* as a function of 〈*N_P_*〉. The influence of different assumed values for 〈*N_r_*〉 over a whole range of possible values is explored in terms of the resulting *K_D_* and *N_max_*. For every value of 〈*N_r_*〉 all six ligand binding curves were calculated anew and fitted with Eq. 13. The graph shows the mean value of the *K_D_* (solid line) and *N_max_* (broken line) relative to those obtained in the limit of (1 + 〈*N_r_*〉)〈*N_r_*〉 → 1. In the table, parameter values for *K_D_* and *N_max_* are listed for two values of 〈*N_r_*〉, which correspond to vesicle diameters of 80 nm and 100 nm, a labeling fraction of 0.004 mol% fluorescent lipid analog and a mean lipid area of *A_L_* = 0.6 nm^2^. The analysis in Fig. 3 is based on a vesicle diameter of 80 nm (〈*N_r_*〉 = 2.5)

## Discussion

To our knowledge, this is the first demonstration of protein-to-liposome binding curves derived from dual-color FCCS. It provides several advantages compared to the single-color FCS approach (8-10):

I. Selectivity: Dual-color FCCS is more specific to the particular binding process of interest than single-color FCS, as both binding partners are labeled. For instance, liposome binding is more easily distinguishable from protein aggregation.
II. Reliability of curve fitting: Determining diffusional amplitudes is generally more straight-forward than determining and interpreting fractions of diffusing particles in two-component fits.
III. Binding model-independence: In contrast to earlier FCS approaches (9), a binding model is not assumed initially, but the data is converted into a standard type of binding curve (22). Ligand binding curves generally provide freedom for conducting subsequent binding analysis with a model of choice.

Here, the simple Langmuir model of independent, equal affinity binding sites was chosen. In this case, in principle, a single FCCS measurement along with a partial titration curve is sufficient to obtain the *K_D_* according to 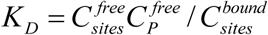, because considering 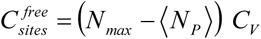 and 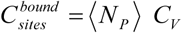 yields

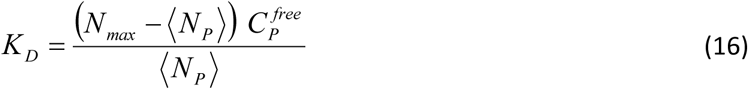

The single titration point consisting of 〈*N_P_*〉 and 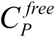 needs to be away from binding saturation to obtain a meaningful value for *N_max_* –〈*N_r_*〉. In addition, *N_max_* needs to be determined from a partial titration curve that goes to saturation.

We chose not to use this “single point” approach, because fitting a complete binding curve yields a more reliable value for the *K_D_*.

(IV) In contrast to (9), we do not assume a Poisson distribution for the bound protein. The Poisson distribution is appropriate for describing the distribution of the small fraction of fluorescent lipid. However, it is less suitable for the protein distribution, because it fails to describe the saturation of protein binding for the case of a finite number of binding sites on the liposome. In the earlier FCS approach, the Poisson distribution was employed to determine the second moment 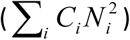 needed for interpreting the autocorrelation curves. Using the cross-correlation amplitude, only the average number of bound proteins is employed, but not the second moment.

(V) Unlike (8, 9), it is not required that liposome morphology is unaffected by protein binding. Indeed, an increase in the diffusion time of the liposomes was observed upon protein binding, which may be partially due to membrane remodeling, but the diffusion times are not needed for the analysis.

In addition, the dual-color FCCS approach shares some advantages with the single-color FCS approach:

I. Sensitivity: Like FCS, dcFCCS requires only minute amounts of samples and short measurement times.
II. Dynamics: Temporal evolution can be monitored and measurements performed at equilibrium. Analysis does not rely on kinetics, such as the commonly used binding analysis by surface plasmon resonance (SPR).
III. Solution measurements: Unlike for example in SPR, liposomes are not immobilized on a support, but are free to diffuse and to adopt native shapes, rather than for instance flattening out on a substrate.

In addition, the dcFCCS data analysis scheme derived here eliminates the need for common preassumptions in dcFCCS analysis:

I. Defined degree of labeling: In dcFCCS, stoichiometric 1:1 labeling of both binding partners is often the goal, because missing labels on bound particles fail to contribute to the cross-correlation. Incomplete green labeling does indeed affect the green auto-correlation amplitude 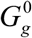 and the ratio of cross-correlation to green autocorrelation amplitude 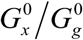. But neither is the absolute cross-correlation amplitude 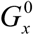 itself altered (due to its normalization with the average green fluorescence) (Eq. 1), nor is the ratio of the cross-correlation to the red auto-correlation amplitude 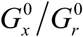 affected. (By the same token, incomplete red labeling affects 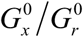, but neither 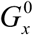 nor 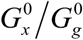 is affected.) Here, we exploited the fact that 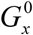 and 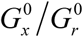 are insensitive to incomplete green labeling and chose to mix green-labeled and unlabeled protein. Because only the mean number of green fluorophores enters into the analysis, it would even be permissible to use multiply labeled ligand molecules (e.g. proteins labeled on lysin residues). The degree of labeling of the red particles in contrast is relevant for 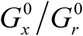 and thus for our binding analysis. Instead of hard-to-achieve stoichiometric labeling, stochastic labeling of liposomes with a Poisson distribution of the red lipid analog was used, which is easily accounted for by the factor 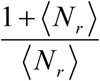 (see Eq. 11).
II. Limited concentration range: By mixing green-labeled and unlabeled protein, suitable numbers of green-labeled particles in the detection volume could be realized over several orders of magnitudes of total protein concentrations. Titrations were performed up to 200 μM of total protein, which is far above suitable concentrations for purely labeled particles (≈ 100 nM in (32)).
III. Limited *K_D_* range: As a consequence of the extended concentration range, not only tight interactions with dissociation constants in the pM to nM range, but weaker interactions in the μM range are accessible by FCCS.

The analysis scheme comes with the disadvantage that the total protein concentration needs to be determined by an independent biochemical method. Because of the proportionality in Eq. 14, the relative error in the protein concentration directly contributes as a relative error to the *K_D_* (Eq. 14). Likewise, the relative error in the protein concentration directly contributes as a relative error to the total number of binding sites *N_max_* (Eq. 13). To minimize the impact of statistical error in the protein concentration determination, each binding experiment was carried out and analyzed independently (Fig. 3) and the *K_D_* values were averaged, yielding 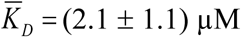..

To our knowledge, the absolute binding affinity of Sar1p to bilayers has been assayed in one other case (33) but under conditions that are not comparable, i.e. a different lipid composition, a labeled nucleotide instead of a labeled protein and non-equilibrium conditions: Loftus et al. quantified the fluorescence intensity of Sar1p incubated with BODIPY FL GTPyS on ‘major’-mix bilayers (lacking the ‘minor’ components such as phosphoinositides) after washing away excess labeled nucleotide and free protein and obtained *K_D_* = (10.5 ± 3.1) μM.

The dual-color FCCS analysis scheme developed here is readily applicable for testing binding models and determining binding affinities under controllable conditions. For example, binding may be studied as a function of protein variants, lipid composition and buffer composition (including nucleotides, salt and pH) to gain insight into specific interactions and into electrostatics. The dcFCCS scheme does not provide a direct view of the structure and the topology of the interaction. Structural biology methods, such as NMR spectroscopy, cryo-electron microscopy and mass spectrometry may be used to obtain complementary insight into the mechanism of the interaction between protein and membrane.

## Conclusion

Performing protein-liposome binding experiments by dcFCCS instead of FCS provides advantages by exploiting amplitudes, eliminating the second moment and leading directly to the degree of protein binding. Moreover, dcFCCS binding analysis is quickly implemented, provided that the optical setup is already capable of dcFCCS, because liposome labeling only involves mixing the lipids with a low percentage of fluorescent lipid. The principle may be extended to any kind of ligand-liposome binding, including ligands binding to liposomes that contain reconstituted proteins, such as membrane receptors. Membrane protein labeling becomes dispensable, because the lipid bilayer can be labeled instead. Binding of non-fluorescent ligands may be measured using a labeled ligand and a competition assay. Furthermore, the principle can be extended to other instances of ligands binding to diffusing particles, including lipid nanodiscs.

## Acknowledgements

We thank Jan Auerswald for fluorescent labeling of Sar1p and Stefan Werner for contributions to the analysis. The BMBF (03Z2HN22), Land Sachsen-Anhalt/ERDF (124112001 and 1241090001) and the DFG (GRK 1026) are acknowledged for funding.

## Author contributions

K.B. designed research, D.K. prepared the samples and performed measurements, D.K. and J.E. analyzed the data, J.E. and K.B. wrote the manuscript.

